# The impact of Agri-Environment Schemes (AES) and red fox (*Vulpes vulpes*) on the density of European brown hare (*Lepus europaeus*) populations in Hungary

**DOI:** 10.1101/2024.06.10.598166

**Authors:** Nikolett Ujhegyi, Zsolt Biró, Veronika Bókony, Norbert Keller, Krisztián Katona, Miklós Heltai, Sándor Csányi, László Szemethy

## Abstract

Most Lepus species can easily adapt to new or changing environments. Until the 1960s, the brown hare estimated stock in Hungary went above 1,200,000 specimens. But also in Hungary and across Europe, hare densities have decreased in the past few decades. The primary cause of declining hare populations is habitat alteration in agricultural landscapes caused by the intensification of agricultural practices. Because the well-being and herd size of small game species, such as the hare, are strongly influenced by the ecological environment, following in their population characteristics can be used as an indicator of habitat quality, with the results being used not only in wildlife management, but also in nature conservation and agricultural greening programs. Our goal is to compare the size of the hare game management to the efficiency of the agricultural support system, vegetation types, and the quantity of red fox, all of which can have an impact on the hare’s density.

In a large-scale survey we found that the estimated hare density index decreased year after year. There were no more animals discovered in the "better areas" overall. According to the model, if the game management units contained a large proportion of AKG arable land, the expected hare density would rise. Similarly, if the region had a high proportion of AKG fields but a low or average fox harvest rate, the hare population would be unable to grow. Animal density decreased slightly, albeit in a trend, as the number of AKG lawns increased. Hare distribution has shrunk over time, especially in areas with a high percentage of suitable habitat. The hare harvest density remained constant year after year in VGEs with a higher proportion of AKG grassland. The fox’s declining rate had no effect on the increase in harvest density. Overall, it appears from a national spatial scale study that even in less good areas, the support could not improve the habitat because its effect did not depend on the proportion of good areas, i.e. even in the case of VGEs with a small proportion of good areas, it could not add much to the hare population density a lot of AKG. Cattle pasture comprised a significant portion of the AKG grasslands. Because the effect of intensively grazed regions on hares is clearly negative due to excessively short grass, the high standard deviation values observed in high proportion AKG grasslands may be explained by this.

## 1. Introduction

During the twentieth century, Europe’s farmland biodiversity declined substantially. With an increasing and urbanizing human population, agricultural productivity will increase while the agroecosystem will decline (Kleijn and Sutherland, 2003; Stoate et al., 2009). As a consequence of intensive agriculture, field size has increased and crop species diversity has decreased, resulting in a homogeneous farmland landscape across Europe (Stoate et al., 2009; Tscharntke et al., 2005). Moreover, the edges of agricultural fields that are important to animals will be narrowed or eliminated (Batáry et al., 2010; Stoate et al., 2009; Tscharntke et al., 2005). It has been estimated that 50% of all species in Europe depend on agricultural habitats, including several endemic and threatened species (Stoate et al., 2009). Agri- environment schemes (AES) were implemented in Europe in the late 1980s to slow the decline of biodiversity. The aim of these programs is that environmentally friendly management can be used as an alternative to organic farming, providing farmland biodiversity while maintaining high yields through ecological intensification (Kovács-Hostyánszki and Báldi, 2012). The major goals of agri-environmental projects are to improve the ecological quality of farmland through soil conservation, biodiversity conservation, and water pollution reduction (Ekroos et al., 2014; Stoate et al., 2009). Mixed farming, or spatial and temporal habitat variability, is essential for birds in arable farmland (Stoate et al., 2009). The diversity of crop types, as well as the proximity of semi-natural habitat patches in arable crops or forest borders, constitute vital ecotones for insects, particularly bees and butterflies, and plants (Stoate et al., 2009). Many national AESs increase landscape heterogenety and thus biodiversity benefits (Stoate et al., 2009; Vasseur et al., 2013), and many studies have been conducted across Europe to study the effects of the AES, however monitoring of AESs varies widely between member states, with levels being insufficient to assess effectiveness in most countries (Critchley et al., 2004; Kleijn and Sutherland, 2003; Stoate et al., 2009).

Birds and insects are the most commonly used indicator species in AES programmes, although the results are frequently inconsistent, and studies have shown that the effectiveness of AES varies between positive and negative (Kleijn and Sutherland, 2003; Zingg et al., 2019). In the United Kingdom, the mosaic landscape of arable lands or high cereal-dominated agricultural fields has a good influence on bird abundance (Stoate et al., 2009); nevertheless, AES habitats must support both overwintering and breeding conditions (McHugh et al., 2017). The efficiency of AES measures in Spain was unclear; it was dependent on landscape variability (Concepción et al., 2020). AES had a favorable overall effect in Finland, but it was insufficient to halt the country’s biodiversity decrease (Stoate et al., 2009). An agri-environmental strategy applied in Poland to conserve the aquatic warbler (*Acrocephalus paludicola*) indicates that AES regions had a favorable effect, whereas other species showed neutral or negative effects. (Budka et al., 2019). In Estonia, increased flower abundance on environmentally friendly management farms had positive significant effect on bumblebee abundance and a marginal benefit to bird species (Marja et al., 2014). The crop type is also determinative for the species (Stoate et al., 2009). Many studies suggested that set-aside land improves landscape variability and aids in the restoration of mosaics typical of mixed agriculture (Stoate et al., 2009; Vasseur et al., 2013).

Traditional practiced grasslands are important habitats for biodiversity in Europe. Ecological issues are largely connected with the more productive grassland forms, as they are with arable systems. Silage management has had a negative influence on bird food supplies, resulting in increased mortality of ground-nesting birds. Permanent grasslands are uncommon in intensive agriculture, with all grasslands being part of crop cycles. Plants, invertebrates, and birds have different grassland management requirements. High density cattle grazing had the potentially negative effect on invertebrate and avian species (Stoate et al., 2009).

In Hungary, a pilot AES was implemented in 1999 and was expanded to the national level in 2004 (Stoate et al., 2009). The most recent revision of the Common Agricultural Policy (CAP) comprised payment decoupling from production, the implementation of cross- compliance, modulation, and the extension of the Rural Development Program. Previous research found that intensifying cereal management with crops had a negative impact on the species richness of birds, bees, and spiders, but had a beneficial effect on the diversity and abundance of carabid beetles (Báldi et al., 2005; Batáry et al., 2015; Stoate et al., 2009). The grazing intensity had a moderate negative effect on orthopterans and beetles and a strong negative effect on grassland birds (Batáry et al., 2015, 2007). Mowing can also increase predation on birds and mammals (Stoate et al., 2009).

However, the impacts of the AES are often positive for insects, but contrary for bird species. Because large-territory bird species can exploit resources over a much larger area than specialized or narrow-territory species (Marja et al., 2014), there is a need for more focused farm-scale AES and the testing of agri-environmental policies on a variety of species, including the European brown hare (*Lepus europaeus*). The brown hare is closely associated with agricultural areas, which can be used to characterize the agricultural landscape (Panek, 2018). Hare is a medium-sized, high-profitability game species with a small home range area, therefore the animals’ survival is strongly dependent on the vegetation and hiding places of arable land, particularly the margins and edge habitats (Edwards et al., 2000; Panek, 2018; Roedenbeck and Voser, 2008). Although it is an important small game species, its populations have been declining throughout Europe since the 1960s (Edwards et al., 2000; Smith et al., 2005). The loss of crop diversity has been identified as a key variable in the reduction of the brown hare (Stoate et al., 2009), as has an increase in the number of predators, particularly red foxes (*Vulpes vulpes*), as well as the spread of illnesses and road mortality (Edwards et al., 2000; Panek, 2018).

Because the brown hare is a hunted game species, as a result, it has been possible to monitor long-term trends in its population using hunting records, because hunting records were a reasonable technique to report temporal trends in the population (Edwards et al., 2000; Tapper and Parsons, 1984). Brown hare hunting records usually indicate how good the breeding season has been from year to year. The trend of hare hunting records and the trend of estimated hare density in spring could be good indicators for environmental interventions or environmental carrying capacity (Edwards et al., 2000). As a result, we employed hunting bag and hare estimation data in our investigation.

AES has been introduced to mitigate the negative environmental effects of intensive agricultural landscapes (Ekroos et al., 2014), they are key tools in the European Union’s efforts to reverse long-term declines in farmland biodiversity (Wilson et al., 2007). However, using AES in extremely complex landscapes with existing high in-field biodiversity levels will provide no extra biodiversity benefits (Concepción et al., 2020). That is why we need to know where we can expect the AES to have an impact. Because mixed farm areas are important to the hare (Smith et al., 2005; Tapper and Barnes, 1986), areas with a high proportion of arable land, various crops, fallow habitats, hedgerows, pastures, woodlands, or shrub increasing the survivor of the hares and leverets (Edwards et al., 2000; Smith et al., 2005; Zellweger-Fischer et al., 2011), because those habitats have higher feeding and sheltering places, so the hares’ mortality rate from predation may be lower (Marboutin and Aebischer, 1996; Smith et al., 2005).

Many studies found that the impact of brown hare predators were different. If the habitat is poor, the fox population may have a significant negative impact on the hare population (Schmidt et al., 2004; Smedshaug et al., 1999; Smith et al., 2005); particularly, changes in fox abundance affected the situation of hares primarily in the autumn-winter season (Misiorowska and Wasilewski, 2012; Panek et al., 2006); however, habitat improvement may be much more effective in restoring prey populations than fox control (Knauer et al., 2010); additionally, many studies find no effect of fox density on the abundance or decline rate of hares (Panek, 2018).

We aimed to find out, using the brown hare as an indicator species, how the objectives of the agri-environmental measure are met, and whether any overall effect can be demonstrated on a large scale, using a low-cost method based on available data to help understand the effect of AES on our indicator and other species populations. As a result, we examined the relationships between population index and habitat indicators to assess the capacity of the various measures included in AES to support biodiversity across our case study regions. The indices describing the abundance and decline rate of hare populations were calculated using hunting bags and estimate rates. Red fox density was also considered as a potential alternative factor influencing hare populations.

## 2. Materials and methods

### 2.1. Monitoring areas

We analyzed the effects of land use and various agri-environmental scheme options on the brown hare. Biodiversity data were collected as part of the ongoing evaluation of AES conducted within the framework of the "New Hungarian" Rural Development Programme 2007-2013 (Hungarian Ministry of Agriculture and Rural Development, 2015). This survey was part of the AES impact monitoring system, controlled by the National Food Chain Safety Office the conclusions were drawn on the effectiveness of the Agri-environmental scheme programs. The future usability of the 21 difference indicators, and the possibilities of carrying out the monitoring system based on the statistical analysis of the available indicators (Table 1). We analyzed the small game species indicators collected during a large-scale survey in Hungary. Furthermore, we extended our measure beyond the AES areas with data on the percentage of good natural habitat for the brown hare and the population density of one of the hare’s most important predators, the red fox.

**Table 1.** The name of the Hungarian AES categories between 2009-2014. The name with italic font are our chosen subschemes.

| categories | name of the zonal schemes | name of the horizontal schemes |
| --- | --- | --- |
| arable schemes | anti-erosion scheme (wind)<br>anti-erosion scheme (water)<br>*nature conservation purpose farming - Red-footed Falcon*<br>*nature conservation purpose farming - bird / small game*<br>*nature conservation purpose farming - wild goose / crane*<br>*nature conservation purpose farming - great bustard* | *management of traditional homesteads (tanya)*<br>*organic farming*<br>*integrated farming* |
| grassland schemes | *nature conservation purpose grassland establishment*<br>*environmental land use change scheme*<br>*nature conservation purpose farming - habitat management*<br>*nature conservation purpose farming - great bustard* | *organic grassland management*<br>*extensive grassland management* |
| plantation schemes |  | management of traditional orchards<br>organic fruit production<br>integrated fruit production |
| wetland schemes | management of wetlands<br>conservation of arable land into wetland | reed management |

### 2.2. GIS data selection

We selected 482 wildlife management units from 1400 in the Hungarian National Game Management Database, whose primary objective of wildlife management was small game, and who are therefore likely to intensify mesocarnivore hunting (Figure 1). We used the Quantum GIS (QGIS) software (Graser, 2016) to clip the shapefiles of the 482 WMUs from the CORINE Land Cover 2012 raster (Gallego and Peedell, 2001), thus we got the habitat type of our WMU-s. The Hungarian AES program (2009-2014) included 21 different zonal and horizontal schemes (Table 1). During this time, the government provided 14.688 grants to altogether 1.163.663 ha throughout Hungary. We selected 13 schemes that were important and relevant to the brown hare (indeed: the study size was similar, and the regulations were positive for the animals) (Table 2). The National Food Chain Safety Office provided us the categorized shapefiles for these selected Hungarian Agri-environmental schemes based on cultivation branch (grassland or arable land). We used QGIS software to clip the WMU shapefiles from the AES shapefiles, yielding the AES areas for each WMU.

**Figure 1.**
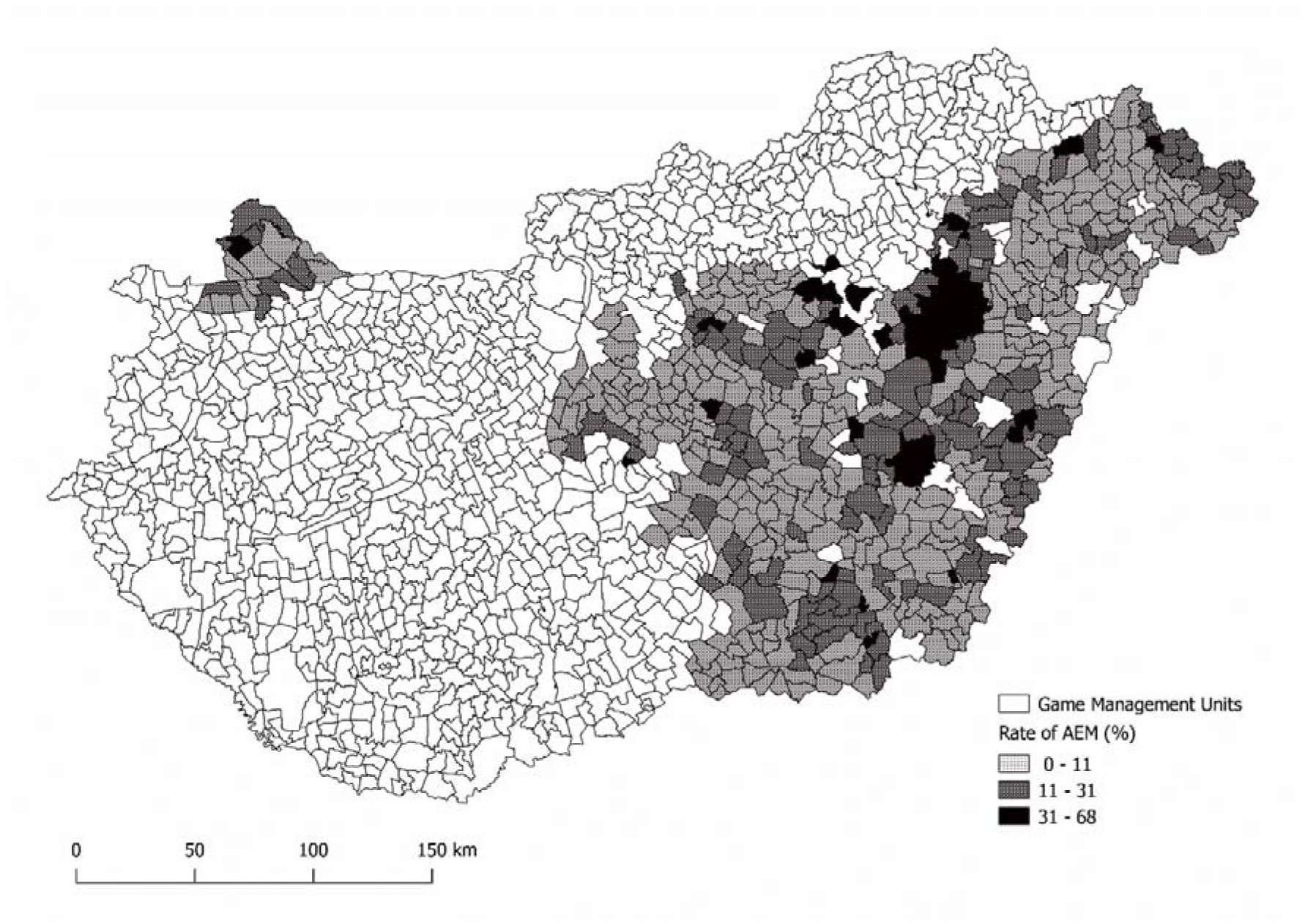
The AEM coverage of Hungary’s chose wildlife management units.

**Table 2.**
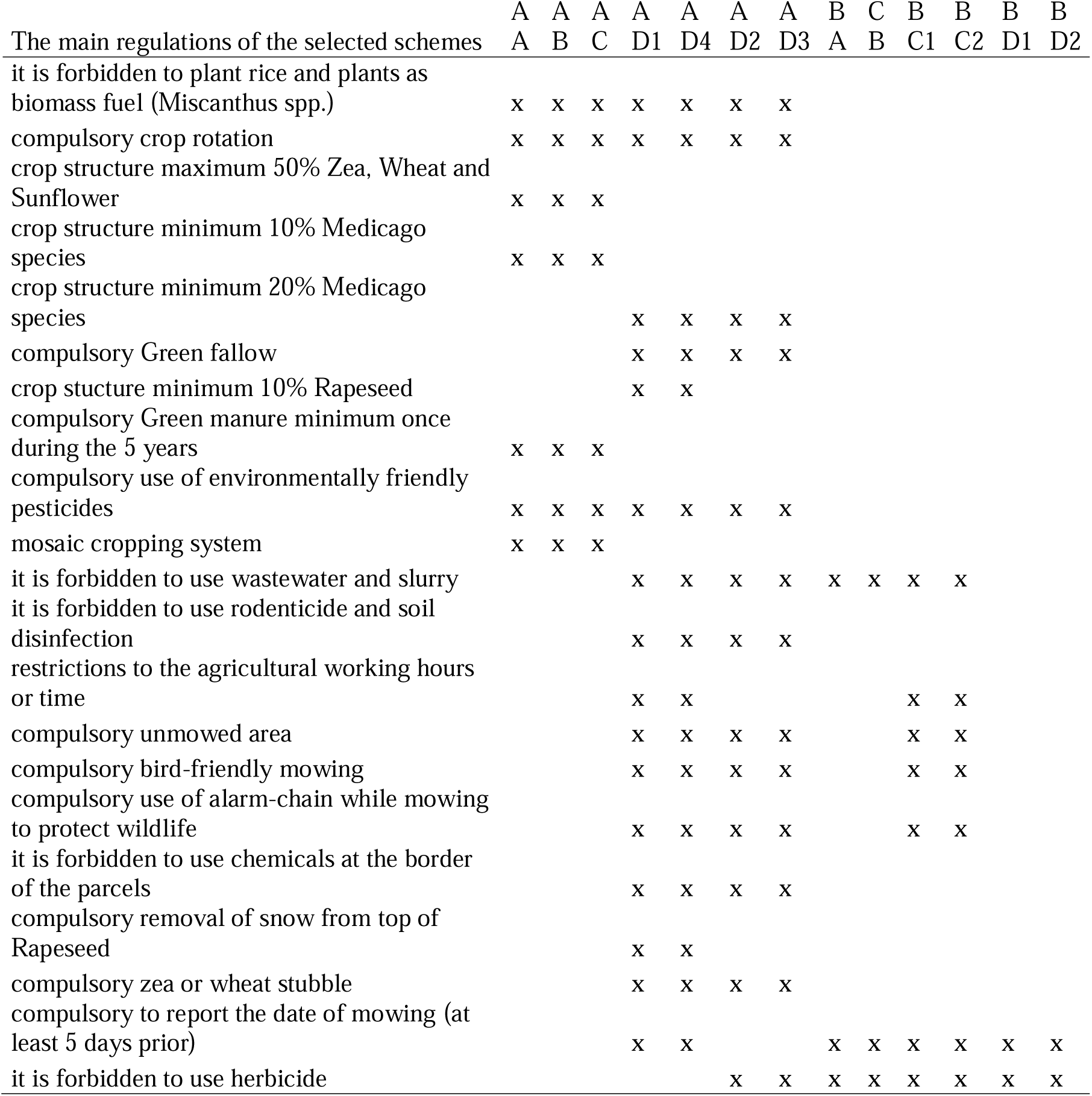
The main criteria of the subschemes, which are included in the study. AA) integrated farming, AB) organic farming, AC) management of traditional homesteads (tanya), AD1) nature conservation purpose farming - great bustard, AD4) nature conservation purpose farming - Red-footed Falcon, AD2) nature conservation purpose farming - wild goose / crane, AD3) nature conservation purpose farming - bird / small game, BA) extensive grassland management, CB) organic grassland management, BC1) nature conservation purpose farming - great bustard, BC2) nature conservation purpose farming - habitat management, BD1) environmental land use change scheme, BD2) nature conservation purpose grassland establishment

### 2.3. Data collection

We selected and categorized these areas, which were preferred habitats for the brown hares based on Pelorosso et al. (2008). The good habitats and the more preferred habitats (altogether 26021.18 km^2^, which were scored 3 and 4) were: non-irrigated arable land, vineyards, pastures, land principally occupied by agriculture, natural grassland, and the other appropriate habitats. The less preferred habitats (altogether 2427.38 km^2^, which were scored 1 and 2) were: fruit trees and berry plantations, complex cultivation patterns, transitional woodland shrubs, and sparsely vegetated areas. These preferred areas were the net areas of the WMU-s (Fig. 2).

**Figure 2.**
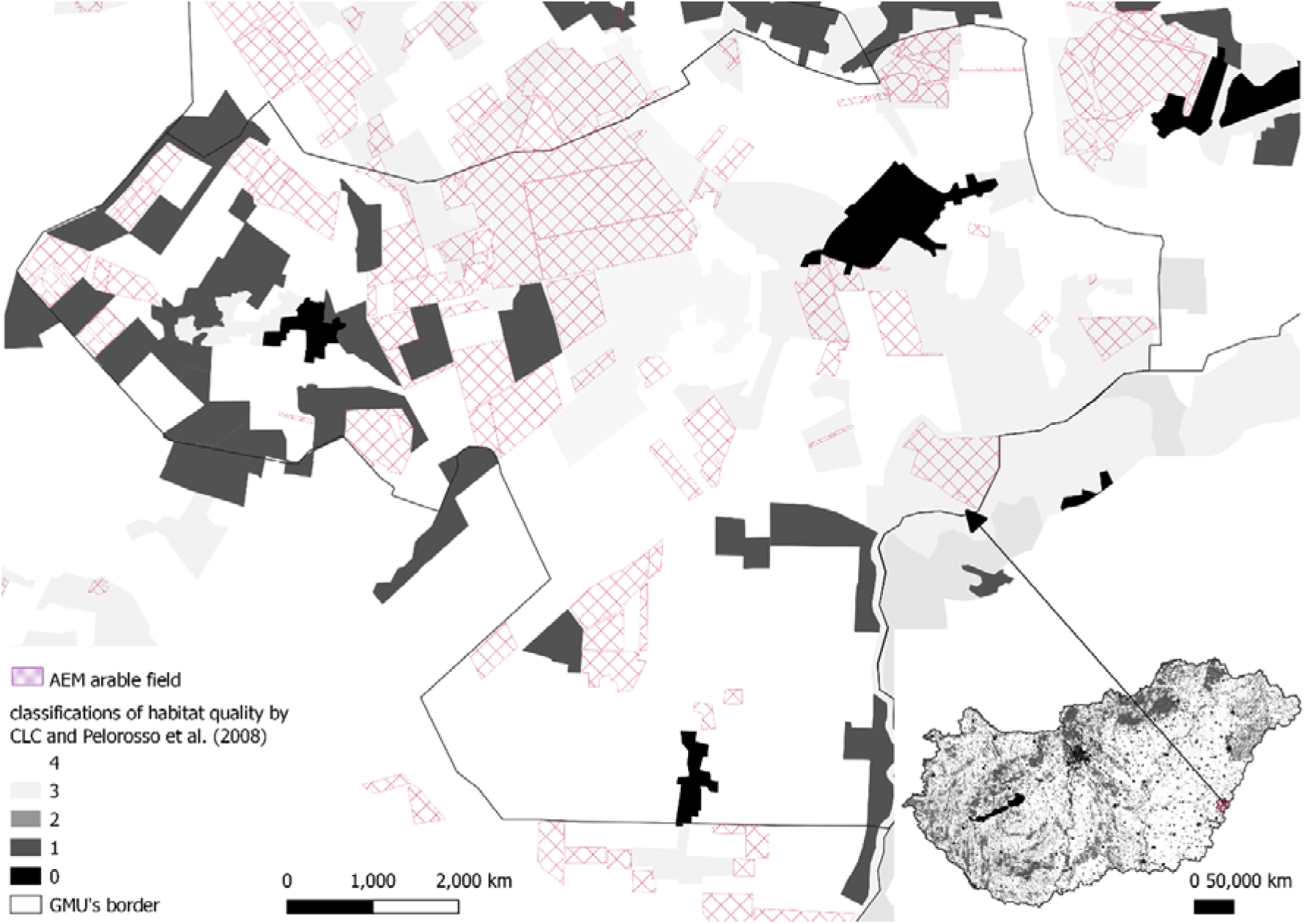
An example of GMU’s CLC area qualities and AEM coverage. The purple areas on the enlarged map detail are AEM areas belonging to one of the 13 target programs, the black spots are unsuitable habitats for rabbits, the places marked 1&2 are less preferred, and the areas marked 3&4 are "good areas".

We calculated the overall and separate AES proportions (%) of the WMUs using the WMU’s net area and two different AES areas. We also calculated these proportions (%) for the WMU’s more preferred areas (good habitat).

We used the Hungarian Game Database’s hare estimate and hunting bag data (2007-2014) to calculate the population and harvest density (individuals/100 ha) of the WMU’s net area. Because this AES began in 2009, we used the mean of the 2008 and 2009 estimate data, as well as the mean of the 2007 and 2008 hunting bag data, as references for the AES. The average of the reference years was subtracted from the given annual population or harvest density.

We selected fox estimates and hunting data from the Game Database for the years 2007 through 2014. We calculated the same data for the fox (except that our rates were individuals/1000 ha). Furthermore, we counted and categorised each WMU’s fox harvest rates. This value was calculated as the harvest density divided by the population density. Then we classified it using Heltai et al. (2010): if it was less than 1.5, the harvest rate was "bad," 1.5–2 was "adequate," and more than 2 was "good."

### 2.4. Statistical analyses

All analyses were run in the R 4.02 computing environment (R Core Team, 2016), using the following packages: nlme (Jose et al., 2018), lattice (Jose et al., 2018), lsmeans (Lenth, 2016), car (Fox et al., 2018), classInt (Bivand et al., 2020). We used linear mixed-effects models (LME) to test whether each hare population index data (the deviation of the population density and harvest density from the initial years) differed across the WMU-s good habitat size, the AES type and size, and the harvest density of the fox according to the fox’s population density size in the former year. Because our main question was the impact of the AES on the hare density, we would like to know whether our model had been multicollinearity among explanatory variables or not. We found that our model "vif" values were less than 2 to the main effects, so we did not find multicollinearity.

We conducted our analyses in two parts. First, we examined the deviation of the hare density from our reference data, which we calculated using the mean of the two years before the AES (2008 and 2009). In this model, we tested whether the brown hare estimate data differed across the WMU’s good habitat size, the AES type (arable land or grassland) and proportion (%), as well as the fox harvest rate and population density in the previous year. Because predators have a functional response effect on their prey (Angerbjörn, 1989; Cosner, 1999), and hare estimation data is collected in the spring, we used fox hunting data from the previous year in this model. Second, we tested the hare harvest density in the same way (because the hunting bag is also a good and useful indicator of this species’ abundance (Panek and Kamieniarz, 1999)), but in this model, we used the same fox data from the previous year because the hunting season is in the autumn and the survived hares are influenced by the foxes of that year.

Our explanatory variables for each model were the logarithmic net area of AES (LOG10) (separately from grassland and arable land), the proportions of good habitats, the years, the density, and the fox harvest rates. We used good habitat, year (as a number), net area of AES arable land, fox density, and fox harvest rate interaction, therefore the WMU identity number was used as a random factor.

We used the same LME model structure for the hare harvest density data, but also included the LOG AES type × good habitat × year interactions (with the year and ID as a random factor) and tested its significance with an analysis-of-deviance test (type-2 ANOVA). Because the AIC was lower in our first model, we only used the ID as a random factor.

Finally, because less WMU had a higher percentage of AES, we classified the "good habitat" and AES data into three groups before running our model with this version, which included a category variable. We also used Jenks’ natural breaks method to divide the values into various natural classes (Gianmarco, 2017).

## 3. Results

### 3.1. The effects of the hare population density rate

We found no relationship between the net AES proportion and the "good habitat" proportion. We found a significant interaction between fox density, fox hunting rate, and the growth rate of the net arable AES area. (Table 3). If the WMUs had a high percentage of AES arable land, the hare population density could increase. However, if the AES arable land was high but the fox harvest rate was weak or adequate, the hare density could not increase.

**Table 3.**
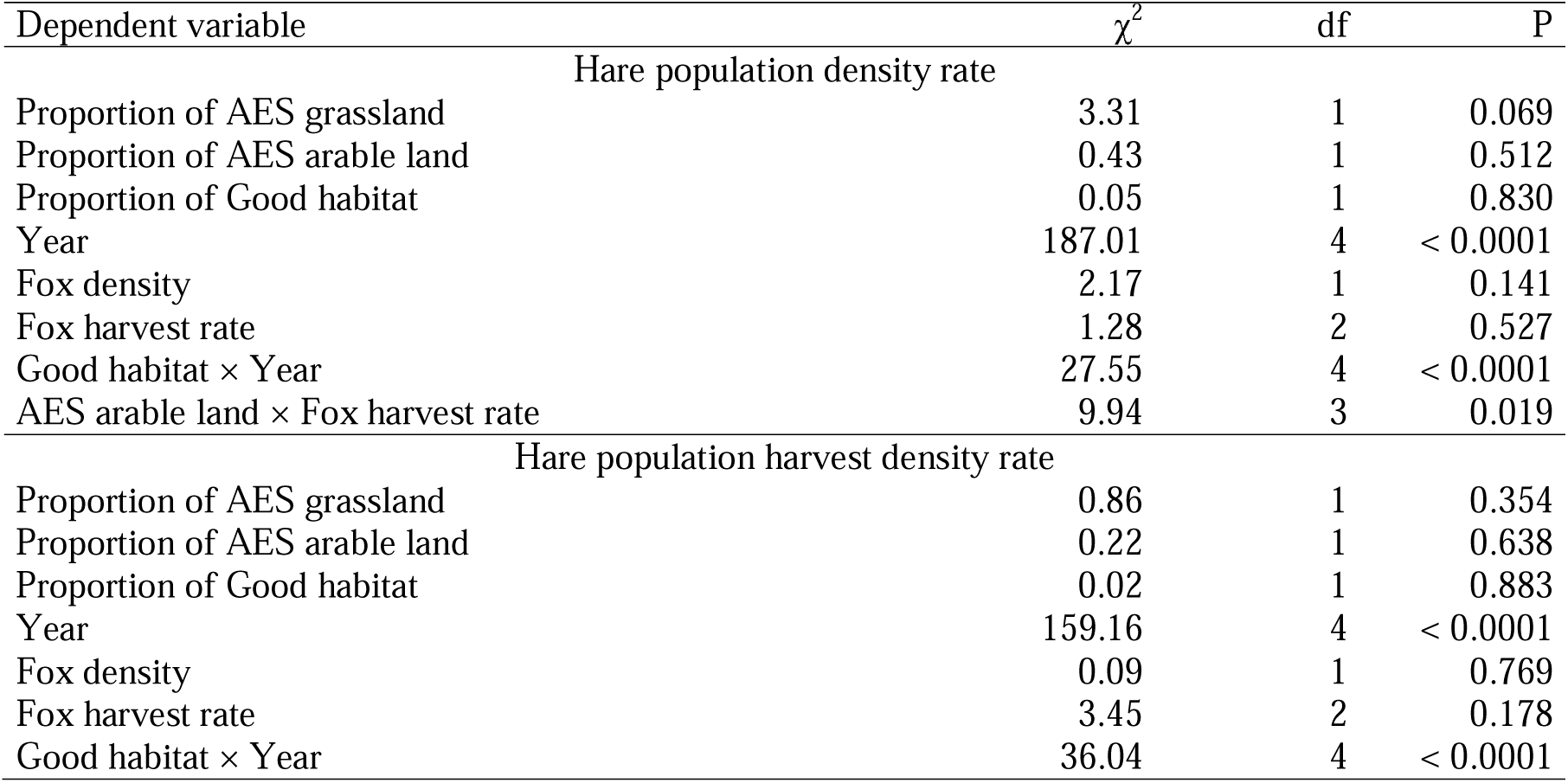
ANOVA tables of the LME models testing the effect of the AES type percent × good habitat percent × year interaction on the two hare density variables, with wildlife management units as random factor.

We found a significant interaction between good habitat percentage and years. The hare density differed from the reference density if the percentage of good habitats increased over the years. The years were also significant. Hare density differed marginally in the AES grassland (Table 3). Higher percentages of AES grassland had a negative impact on hare density (linear contrast: b ± SE = -0.37 ± 0.2, t_478_ = -1.82, P = 0.069).

If we used Jenks’ natural breaks classification method to categorize the good habitat percent, we could determine the relationship between the good habitat and the years. In this model, the year remains significant, and the AES arable land category has a significant impact on the fox harvest rate. Neither the categorized AES arable land, grassland, or good habitat indicated any significant differences (Table 4). 2010 and 2013 were the two worst years. In 2009 the difference from the reference year was higher in the WMU-s, where the good habitat percentage was high, or the AES arable land was over 26%. The high percentage of AES grassland was not favorable for the hare. It seems that the arable AES and the high percentage of good habitat can have a positive effect on the hare population because in 2011 those two categories could have lifted the collapsed hare population (Fig. 3). It seems that the high fox harvest rate cannot exert an effect on its own. In general, the hare population density index decreased over the years, especially in the weakest treatment category. The highest treatment category was able to break the downward trend (Fig 3).

**Figure 3.**
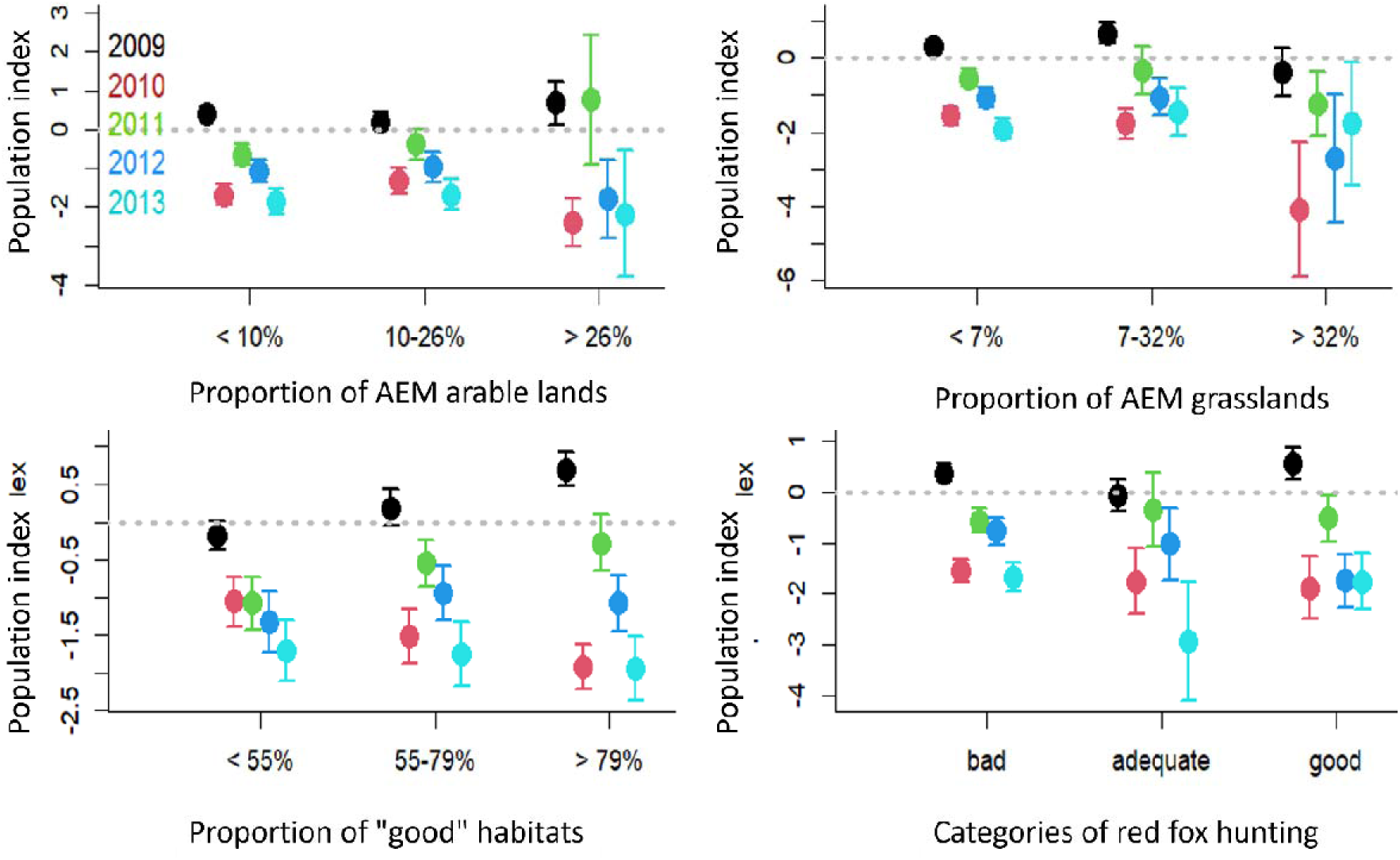
Changes in the coverage of AKG treatments, the percentage of good areas, or the categories of the fox reduction rate influence the variation in the estimated hare density against the reference years. According to estimates made during the reference period, the dashed line at zero represents the stock density.

**Table 4.** ANOVA tables of the LME models testing the effect of the AES type category × good habitat category × year interaction on the two hare density variables, with wildlife management units as random factor.

| Tényezők | $\chi^2$ | df | P |
| --- | --- | --- | --- |
| Hare population density rate |  |  |  |
| AES grassland in 3 categories | 0.6589 | 2 | 0.71933 |
| AES arable land in 3 categories | 0.0736 | 2 | 0.96389 |
| Good habitat in 3 categories | 0.0994 | 2 | 0.95153 |
| Year | 182.0571 | 4 | < 0.0001 |
| Fox density | 2.2178 | 1 | 0.13642 |
| Fox harvest rate | 1.1484 | 2 | 0.56315 |
| Good habitat in 3 categories × Year | 17.7291 | 8 | 0.02335 |
| AES arable land in 3 categories × Fox harvest rate | 27.3377 | 4 | < 0.0001 |
| Hare population harvest density rate |  |  |  |
| AES grassland in 3 categories | 1.1987 | 2 | 0.5492 |
| AES arable land in 3 categories | 2.6358 | 2 | 0.2677 |
| Good habitat in 3 categories | 1.4945 | 2 | 0.4737 |
| Year | 158.6455 | 4 | < 0.0001 |
| Fox density | 0.0385 | 1 | 0.8445 |
| Fox harvest rate | 3.83962 | 2 | 0.1466 |
| Good habitat in 3 categories × Year | 32.0493 | 8 | < 0.0001 |

### 3.2. The effects of hare harvest density rate

We found a significant interaction between the good habitat percentage and the years. The hare harvest density deviation differed from the reference density between years (Table 3). The least hare was shot in 2010, except in that WMU, where the AES grassland had high percent. The categorized habitats have shown significant interaction over the years. And the year was also significant (Table 4). In 2010 the hare harvest density significantly increased (linear contrast: b ± SE = 0.92 ± 0.48, t_1917_ = 1.93, P = 0.053) compared to the reference years, and 2011 had a reducing effect on the harvest density (linear contrast: b ± SE = -0.32 ± 0.48, t_1917_ = -0.68, P = 0.496). In that WMU-s which had higher AES grasslands the hare harvest density was similar every year (Fig 4). In the two bad years (2011 and 2013) the hare harvest density was also lower (except for the WMU-s with high grassland AES).

**Figure 4.**
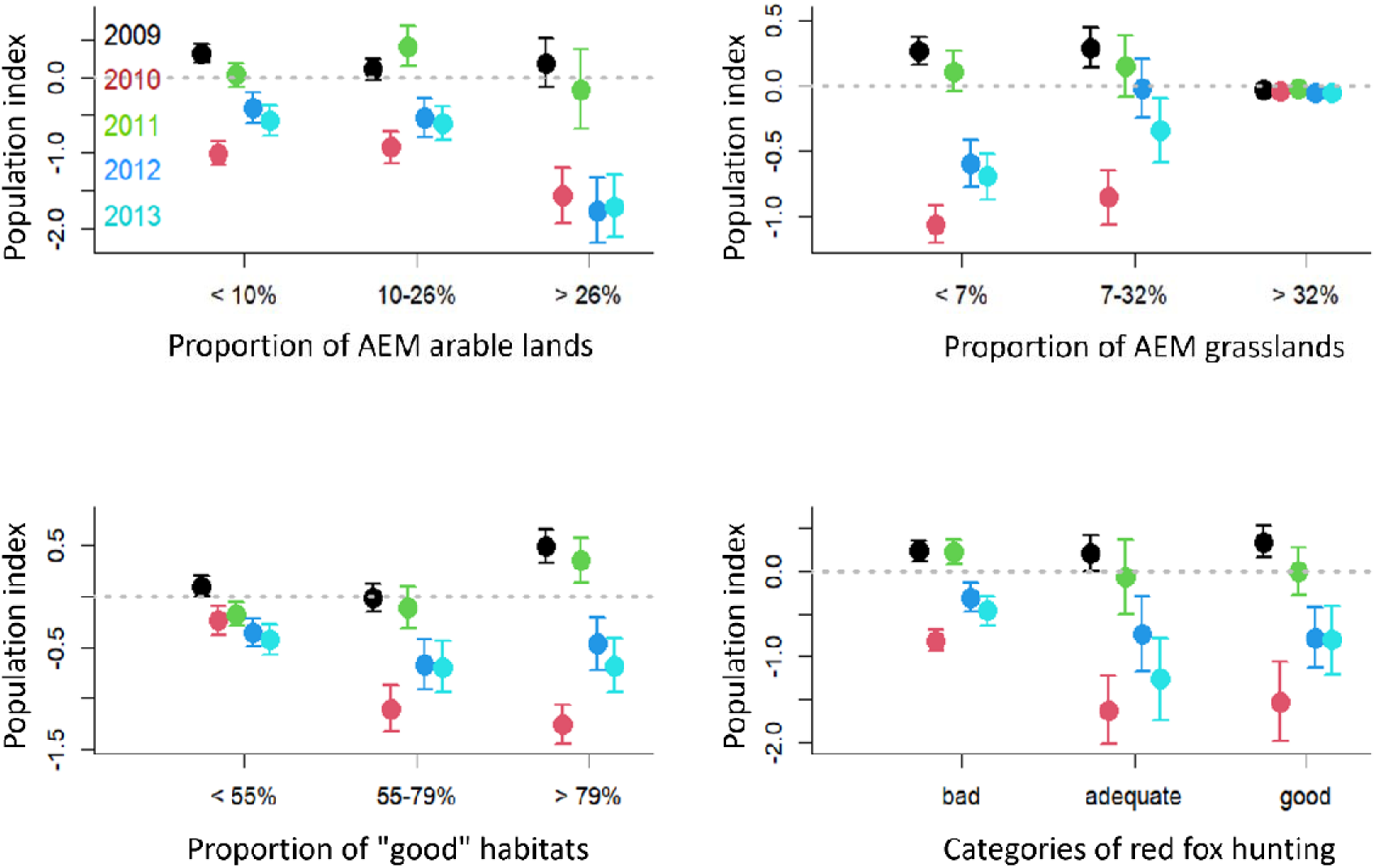
Changes in the coverage of AKG treatments, the percentage of good areas, or the categories of the fox reduction rate influence the variation in the estimated hare harvest density against the reference years. According to estimates made during the reference period, the dashed line at zero represents the stock density.

## 4. Conclusions

It seems the AES in Hungary did not depend on whether the arable land contained more good habitat or not. Because the main aim of the AES is habitat improvement if it is applied to that area, which had a mosaic habitat structure, and a lot of feeding and hiding places, the AES could not exert a positive effect. AES’s larger percentage of arable land may be affected by hare density; if the year’s weather is hare-friendly, the high percentage of arable land could prevent the hare population decline. At the same time in Hungary in that AES period between 2009-2014, the highest AEM arable land proportion of the WMU-s was 60 %. In some years it may indicate some positive effects on brown hare populations such as in (Zellweger-Fischer et al., 2011)s study, but the quantity and quality of AES must still be increased. Because the vegetation quantity of the field is important for the hare (Langhammer et al., 2017; Zellweger- Fischer et al., 2011), the crop fields of large acreage have a negative impact on hares (Panek and Kamieniarz, 1999), in the future we need to produce more permanent set-aside sites and herbaceous field margins, because these interventions have the largest population response effects (Langhammer et al., 2017). We need to know the fields’ vegetation in each year and season during the AES in the future, because when the WMUs had higher percentage from the avoided vegetation, the potential positive effect of AES could not prevail (Ujhegyi et al., 2021). In this Hungarian AES the maximum size of each parcel was 75 ha, but if there was not preferred vegetation for the hare on the most parcels, that did not help for the hare to survive. The actual vegetation of each treated parcel is needed to know or assess the impact of the AES in small scale survey.

The most of the AES grassland was cattle pasture. The grass out of rotation tended to be positively affect (Schmidt et al., 2004), but heavily grazed areas had a negative effect to the hare (Fourcade et al., 2018), because the grass is too short in that area and the hares have no hiding places. We found a high standard deviation in the high category of AES grassland, suggesting that vegetation height may be more significant than whether a grassland parcel is treated with AES or not. Therefore, we require finer scale in grassland areas.

If a habitat had mixture of crop types, it can provide habitats for species of different ecological profiles (Stoate et al., 2009). The "Good habitat" category could not show unequivocal positive effect in every year (however the standard errors were smaller if the good habitat was high compared to AES arable land), but the most part of that category was arable land. That is similar to the AES arable land if in some years the most vegetation type of the fields was crop plant (especially in large area), we could not see the positive effect to the hare. Because the wild flowers, hedge and field margins are very important not just for the brown hare (Alison et al., 2017; Fischer and Wagner, 2016), in the future we need to know which parcel have flowering meadow strip, or uncutted field borders, which will be feeding and sheltering habitat to the hare, that could increase the survival rate of the animals. This is why the AES supported area must be chosen carefully (Ausden and Hirons, 2002).

Sometimes we could see the positive impact of the agri-environment treatment, if it has been working long enough time (Lindenmayer et al., 2012). For example, in Portugal, positive population trends for the great bustard have been detected where agri-environment measures targeted at steppe birds have been in place for at least a decade (Stoate et al., 2009). It is possible on the reference year the WMUs had also AES support from the previous AES program.

It seems the red fox density or harvest rate could not affect to the hare abundance, however many studies showed that the fox had an impact on the hare abundance (Panek et al., 2006) but in some studies changes in agricultural landscapes seemed to have an even stronger impact on hare density than red fox predation (Knauer et al., 2010; Schmidt et al., 2004), so if the habitat is not appropriate to the hare, the high harvest rate of fox cannot help to increase the hare population. Probably the good habitat or a milder weather have a beneficial effect to the fox density too. In 2010 the weather in Hungary was full of extreme records. In the first quarter of the year snowed nationwide, at April and May, when is a peek of the breeding season (Flux, 2009; Lincoln, 1974) for the hare was four times more rain, than average. In this year Hungary had a record amount of rain with stormy winds with cold fronts and inland waters (Móring and Kolláth, 2011). In 2013 the weather was also full of extreme records. Thunderstorms in winter, snowstorm in March, big flood on the Danube in June, heatwave in mid-summer (Fodor et al., 2014).

Other subsidies combined with the AES could explain enough effect to the hare population growing. The CAP reform encourages field margins, trees, and the prevention of field size expansion. Therefore, when paired with voluntary metrics that are more focused on the local area, performance evaluation and adaptation of their ecological success may be identified, possibly even during harsh weather (Concepción et al., 2020). In the future it will be a good opportunity to monitor other greening measures and AES impacts together. Therefore, measuring other agricultural landscapes (e.g., field margins, hedgerows, fallow lands, pastures or woodlands) in small scale survey would be important.

## Acknowledgements

We are very grateful to Gergely Schally for help with data collecting. This work was supported by the "NÉBIH" (04.2/6342-1/2013) and Szent István University "Research Centre of Excellence-9878/2015/FEKUT" project. The project is co-financed by the European Union and the European Social Fund.

Supplementary material

## Author contributions

Hare and fox estimate and hunting bag data were collected and given by SCs, GIS data were collected by NU. Planning analyses by ZsB, KK, MH, LSz, NK. Statistical analyses by VB, NU and ZsB. The first draft was written by NU; all authors contributed to finalizing the manuscript.

## References

1. Alison, J., Duffield, S.J., Morecroft, M.D., Marrs, R.H., Hodgson, J.A., 2017. Successful restoration of moth abundance and species-richness in grassland created under agri- environment schemes. Biol Conserv 213, 51–58. 10.1016/j.biocon.2017.07.003

2. Angerbjörn, A., 1989. Mountain hare populations on islands: effects of predation by red fox. Oecologia 81, 335–340.

3. Ausden, M., Hirons, G.J.M., 2002. Grassland nature reserves for breeding wading birds in England and the implications for the ESA agri-environment scheme. Biol Conserv 106, 279–291. 10.1016/S0006-3207(01)00254-3

4. Báldi, A., Batáry, P., Erdos, S., 2005. Effects of grazing intensity on bird assemblages and populations of Hungarian grasslands. Agric Ecosyst Environ 108, 251–263. 10.1016/j.agee.2005.02.006

5. Batáry, P., Báldi, A., Szél, G., Podlussány, A., Rozner, I., Erdos, S., 2007. Responses of grassland specialist and generalist beetles to management and landscape complexity: Biodiversity research. Divers Distrib 13, 196–202. 10.1111/j.1472-4642.2006.00309.x

6. Batáry, P., Dicks, L. v., Kleijn, D., Sutherland, W.J., 2015. The role of agri-environment schemes in conservation and environmental management. Conservation Biology 29, 1006–1016. 10.1111/cobi.12536

7. Batáry, P., Matthiesen, T., Tscharntke, T., 2010. Landscape-moderated importance of hedges in conserving farmland bird diversity of organic vs. conventional croplands and grasslands. Biol Conserv 143, 2020–2027. 10.1016/j.biocon.2010.05.005

8. Bivand, R., Ono, H., Dunlap, R., Stigler, M., Denney, B., Hernangómez, D., 2020. Package ‘ classInt.’

9. Budka, M., Jobda, M., Szałański, P., Piórkowski, H., 2019. Effect of agri-environment measure for the aquatic warbler on bird biodiversity in the extensively managed landscape of Biebrza Marshes (Poland). Biol Conserv 239, 108279. 10.1016/j.biocon.2019.108279

10. Concepción, E.D., Aneva, I., Jay, M., Lukanov, S., Marsden, K., Moreno, G., Oppermann, R., Pardo, A., Piskol, S., Rolo, V., Schraml, A., Díaz, M., 2020. Optimizing biodiversity gain of European agriculture through regional targeting and adaptive management of conservation tools. Biol Conserv 241, 108384. 10.1016/j.biocon.2019.108384

11. Cosner, C., 1999. Effects of Spatial Grouping on the Functional Response of Predators. Theor Popul Biol 56, 65–75.

12. Critchley, C.N.R., Allen, D.S., Fowbert, J.A., Mole, A.C., Gundrey, A.L., 2004. Habitat establishment on arable land: Assessment of an agri-environment scheme in England, UK. Biol Conserv 119, 429–442. 10.1016/j.biocon.2004.01.004

13. Edwards, P.J., Fletcher, M.R., Berny, P., 2000. Review of the factors affecting the decline of the European brown hare, Lepus europaeus (Pallas; 1778) and the use of wildlife incident data to evaluate the significance of paraquat. Agric Ecosyst Environ 79, 95–103. 10.1016/S0167-8809(99)00153-X

14. Ekroos, J., Olsson, O., Rundlöf, M., Wätzold, F., Smith, H.G., 2014. Optimizing agri- environment schemes for biodiversity, ecosystem services or both? Biol Conserv 172, 65–71. 10.1016/j.biocon.2014.02.013

15. Fischer, C., Wagner, C., 2016. Can agri-environmental schemes enhance non-target species? Effects of sown wildflower fields on the common hamster (Cricetus cricetus) at local and landscape scales. Biol Conserv 194, 168–175. 10.1016/j.biocon.2015.12.021

16. Flux, J.E.C., 2009. Timing of the breeding season in the hare, Lepus europaeus Pallas, and rabbit, Oryctolagus cuniculus (L.). Mammalia 29, 557–562.

17. Fodor, Z., Kolláth, K., Csonka, T., Véber, I., Vincze, E., 2014. Beszámoló 2013 . év éghajlatáról és szélsőséges időjárási eseményeiről.

18. Fourcade, Y., Besnard, A.G., Beslot, E., Hennique, S., Mourgaud, G., Berdin, G., Secondi, J., 2018. Habitat selection in a dynamic seasonal environment: Vegetation composition drives the choice of the breeding habitat for the community of passerines in floodplain grasslands. Biol Conserv 228, 301–309. 10.1016/j.biocon.2018.11.007

19. Fox, J., Weisberg, S., Price, B., Adler, D., Bates, D., Baud-Bovy, G., Bolker, B., Ellison, S., Firth, D., Friendly, M., Gorjanc, G., Graves, S., Heiberger, R., Laboissiere, R., Maechler, M., Monette, G., Murdoch, D., Nilsson, H., Ogle, D., Ripley, B., Venables, W., Walker, S., Winsemius, D., Zeileis, A., R-Core, 2018. Companion to applied regression [WWW Document]. URL https://cran.r-project.org/web/packages/car/citation.html

20. Gallego, J., Peedell, S.i, 2001. Using CORINE Land Cover to map population density, Topic report 6/2001. Copenhagen.

21. Gianmarco, A., 2017. “plotJenks”: R function for plotting univariate classification using Jenks’ natural break method. 10.13140/RG.2.2.18011.05929

22. Graser, A., 2016. Learning QGIS Third Edition, Packt Pubf. ed. Bfirmfingham. Hungarian Ministry of Agriculture and Rural Development, 2015. “New Hungary” Rural Development Programme 2007-2013 1–552.

23. Jose, P., Douglas, B., Saikat, D., Deepayan, S., 2018. Package “nlme”: Linear and nonlinear mixed effects models [WWW Document]. Team, R Core. URL https://cran.r-project.org/package=nlme

24. Kleijn, D., Sutherland, W.J., 2003. How effective are European agri-environment schemes in conserving and promoting biodiversity? Journal of Applied Ecology 40, 947–969. 10.1111/j.1365-2664.2003.00868.x

25. Knauer, F., Küchenhoff, H., Pilz, S., 2010. A statistical analysis of the relationship between red fox Vulpes vulpes and its prey species (grey partridge Perdix perdix, brown hare Lepus europaeus and rabbit Oryctolagus cuniculus) in Western Germany from 1958 to 1998. Wildlife Biol 16, 56–65. 10.2981/07-040

26. Kovács-Hostyánszki, A., Báldi, A., 2012. Set-aside fields in agri-environment schemes can replace the market-driven abolishment of fallows. Biol Conserv 152, 196–203. 10.1016/j.biocon.2012.03.039

27. Langhammer, M., Grimm, V., Pütz, S., Topping, C.J., 2017. A modelling approach to evaluating the effectiveness of Ecological Focus Areas: The case of the European brown hare. Land use policy 61, 63–79. 10.1016/j.landusepol.2016.11.004

28. Lenth, R. V., 2016. Least-Squares Means: The R Package lsmeans. J Stat Softw 69, 1–33. 10.18637/jss.v069.i01

29. Lincoln, G.A., 1974. Reproduction and “March madness” in the Brown hare, Lepus europaeus. J Zool 174, 1–14.

30. Lindenmayer, D., Wood, J., Montague-Drake, R., Michael, D., Crane, M., Okada, S., MacGregor, C., Gibbons, P., 2012. Is biodiversity management effective? Cross- sectional relationships between management, bird response and vegetation attributes in an Australian agri-environment scheme. Biol Conserv 152, 62–73. 10.1016/j.biocon.2012.02.026

31. Marboutin, E., Aebischer, N.J., 1996. Does harvesting arable crops influence the behaviour of the European hare (Lepus europaeus)? Wildlife Biol 2, 83–91. 10.2981/wlb.1996.036

32. Marja, R., Herzon, I., Viik, E., Elts, J., Mänd, M., Tscharntke, T., Batáry, P., 2014. Environmentally friendly management as an intermediate strategy between organic and conventional agriculture to support biodiversity. Biol Conserv 178, 146–154. 10.1016/j.biocon.2014.08.005

33. McHugh, N.M., Prior, M., Grice, P. V., Leather, S.R., Holland, J.M., 2017. Agri- environmental measures and the breeding ecology of a declining farmland bird. Biol Conserv 212, 230–239. 10.1016/j.biocon.2017.06.023

34. Misiorowska, M., Wasilewski, M., 2012. Survival and causes of death among released brown hares (Lepus europaeus Pallas, 1778) in Central Poland. Acta Theriol (Warsz) 57, 305–312. 10.1007/s13364-012-0081-1

35. Móring, A., Kolláth, K., 2011. Beszámoló 2010 . év éghajlatáról és szélsőséges időjárási eseményeiről.

36. Panek, M., 2018. Habitat factors associated with the decline in brown hare abundance in Poland in the beginning of the 21st century. Ecol Indic 85, 915–920. 10.1016/j.ecolind.2017.11.036

37. Panek, M., Kamieniarz, R., 1999. Relationships between density of brown hare Lepus europaeus and landscape structure in Poland in the 1981-1995. Acta Theriol (Warsz) 44, 67–75.

38. Panek, M., Kamieniarz, R., Bresiński, W., 2006. The effect of experimental removal of red foxes Vulpes vulpes on spring density of brown hares Lepus europaeus in western Poland. Acta Theriol (Warsz) 51, 187–193. 10.1007/BF03192670

39. Pelorosso, R., Boccia, L., Amici, A., 2008. Simulating Brown hare (Lepus europaeus Pallas) dispersion: A tool for wildlife management of wide areas. Ital J Anim Sci 7, 335–350. 10.4081/ijas.2008.335

40. R Core Team, 2016. R: A language and environment for statistical computing. R Foundation for Statistical Computing V, Austria. 2016 [WWW Document]. URL https://www.r-project.org/

41. Roedenbeck, I.A., Voser, P., 2008. Effects of roads on spatial distribution, abundance and mortality of brown hare (Lepus europaeus) in Switzerland. Eur J Wildl Res 54, 425–437. 10.1007/s10344-007-0166-3

42. Schmidt, N.M., Asferg, T., Forchhammer, M.C., 2004. Long-term patterns in European brown hare population dynamics in Denmark: Effects of agriculture, predation and climate. BMC Ecol 4, 1–7. 10.1186/1472-6785-4-15

43. Smedshaug, C.A., Selås, V., Lund, S.E., Sonerud, G.A., 1999. The effect of a natural reduction of red fox Vulpes vulpes on small game hunting bags in Norway. Wildlife Biol 5, 157. 10.2981/wlb.1999.020

44. Smith, R.K., Jennings, N. V., Harris, S., 2005. A quantitative analysis of the abundance and demography of European hares Lepus europaeus in relation to habitat type, intensity of agriculture and climate. Mamm Rev 35, 1–24. 10.1111/j.1365-2907.2005.00057.x

45. Stoate, C., Báldi, A., Beja, P., Boatman, N.D., Herzon, I., van Doorn, A., de Snoo, G.R., Rakosy, L., Ramwell, C., 2009. Ecological impacts of early 21st century agricultural change in Europe - A review. J Environ Manage 91, 22–46. 10.1016/j.jenvman.2009.07.005

46. Tapper, S., Parsons, N., 1984. The changing status of the brown hare (Lepus europaeus) in Britain. Mamm Rev 14, 57–70.

47. Tapper, S.C., Barnes, R.F.W., 1986. Influence of Farming Practice on the Ecology of the Brown Hare (Lepus europaeus). J Appl Ecol 23, 39. 10.2307/2403079

48. Tscharntke, T., Klein, A.M., Kruess, A., Steffan-Dewenter, I., Thies, C., 2005. Landscape perspectives on agricultural intensification and biodiversity - Ecosystem service management. Ecol Lett 8, 857–874. 10.1111/j.1461-0248.2005.00782.x

49. Ujhegyi, N., Keller, N., Patkó, L., Biró, Z., Tóth, B., Szemethy, L., 2021. Agri-Environment Schemes Do Not Support Brown Hare Populations Due To Inadequate Scheme Application. Acta Zoologica Academiae Scientiarum Hungaricae 67, 263–288. 10.17109/AZH.67.3.263.2021

50. Vasseur, C., Joannon, A., Aviron, S., Burel, F., Meynard, J.M., Baudry, J., 2013. The cropping systems mosaic: How does the hidden heterogeneity of agricultural landscapes drive arthropod populations? Agric Ecosyst Environ 166, 3–14. 10.1016/j.agee.2012.08.013

51. Wilson, A., Vickery, J., Pendlebury, C., 2007. Agri-environment schemes as a tool for reversing declining populations of grassland waders: Mixed benefits from Environmentally Sensitive Areas in England. Biol Conserv 136, 128–135. 10.1016/j.biocon.2006.11.010

52. Zellweger-Fischer, J., Kéry, M., Pasinelli, G., 2011. Population trends of brown hares in Switzerland: The role of land-use and ecological compensation areas. Biol Conserv 144, 1364–1373. 10.1016/j.biocon.2010.11.021

53. Zingg, S., Ritschard, E., Arlettaz, R., Humbert, J.Y., 2019. Increasing the proportion and quality of land under agri-environment schemes promotes birds and butterflies at the landscape scale. Biol Conserv 231, 39–48. 10.1016/j.biocon.2018.12.022

